# Sub-atomic resolution X-ray diffraction of the SH3 domain from the post-synaptic density protein Shank3

**DOI:** 10.1101/051425

**Authors:** Srinivas Kumar Ponna, Matti Myllykoski, Tobias M. Boeckers, Petri Kursula

## Abstract

The post-synaptic density multidomain scaffolding proteins of the Shank family are structurally poorly characterised. The Shank family consists of three members, and domain-specific interactions of Shank are involved in forming a network of proteins at the post-synaptic region for intracellular signalling and cellular scaffolding. While X-ray crystallography has provided some information on individual Shank domains, the structural basis of Shank interactions is still largely unknown. In this study, the production and crystallisation of the previously uncharacterised Shank3 SH3 domain is presented. The highly twinned crystals diffracted synchrotron X-rays to a resolution higher than 0.9 Å, and these crystals will eventually have the potential to provide an ultrahigh-resolution view into the Shank family SH3 domains and their interactions.

## Introduction

The Shank proteins were first identified by different research groups in 1999, as novel proteins expressed in human and rodent brains, and being concentrated at the postsynaptic density (Boeckers *et al.*, 1999; Naisbitt *et al.*, 1999; Tu *et al.*, 1999; Zitzer *et al.*, 1999). Shank family proteins (Shank1, Shank2, and Shank3) are postsynaptic density (PSD) scaffolding proteins. They are multidomain proteins, which coordinate protein-protein interactions at the PSD, and they can interact directly or indirectly with postsynaptic receptors, such as NMDA type and metabotropic glutamate receptors, as well as with the actin-based cytoskeleton (Boeckers *et al.*, 2002). Full-length Shank proteins consist of an N-terminal region, multiple ankyrin repeats, a Src homology 3 domain (SH3), a PDZ domain, a long proline-rich region, and a sterile alpha motif (SAM). All these domains are believed to have specific interactions with partner proteins for intracellular signalling. The binding partners for the Shank3 SH3 (Shank3-SH3) domain are currently unknown for all the Shank proteins; a possible interaction between Shank3-SH3 and phospholipase Cβ1b, which carries proline-rich segments in its C terminus, was suggested (Grubb *et al.*, 2011), but such a direct interaction has not been confirmed at the molecular level. Another suggested SH3 interaction was reported for GRIP1 (Sheng & Kim, 2000), but also that interaction has remained unproven at the molecular level.

In the brain, Shank proteins are mainly expressed in the cortex, hippocampus, and amygdala and moderately in thalamus and substantia nigra; neuron-specific expression is seen in the cerebellum (Boeckers *et al.*, 1999; Zitzer *et al.*, 1999). Shank3 is one of the most heavily studied proteins involved in neurological disorders, and mutations in Shank are implicated in many neurological diseases. For example, a *de novo* mutation in Shank3 was reported to cause schizophrenia (Grabrucker *et al.*, 2014). Mutations in the Shank3 gene have been reported to cause autism spectrum disorders (ASD) (Durand *et al.*, 2006). On the other hand, overexpression of Shank3 causes hyperkinetic neuropsycopathy disorders (Han *et al.*, 2013). All these disorders are likely to be related to disturbances in the interactions at the postsynaptic density.

Understanding the nature of a protein at the molecular level will give a detailed insight into disorders caused by mutations. Since Shank3 has domain-specific interactions at the post synapse, understanding the structural and functional aspects of different domains will give detailed information about the behaviour of the domain and its interactions with partner proteins. The SH3 domains of the three different Shank proteins are very similar; on the other hand, their sequence is poorly conserved compared to other SH3 domains. Hence, a structural study can be expected to shed light on Shank-specific features and functions of SH3 domains. In this paper, we present protein production, crystallisation conditions, and diffraction data collection for the Shank3 SH3 domain. The crystals diffracting to sub-atomic resolution turned out to be near-perfectly twinned.

## Materials & Methods

The region encoding the SH3 domain of rat Shank3 (residues 470-528; Table 1) was amplified by PCR, using rat Shank3 cDNA as template, and subcloned into the pDONR221 vector. The Gateway system, based on homologous recombination, was used to further transfer the insert into pTH27 (Hammarström *et al.*, 2006).

**Table 1.**
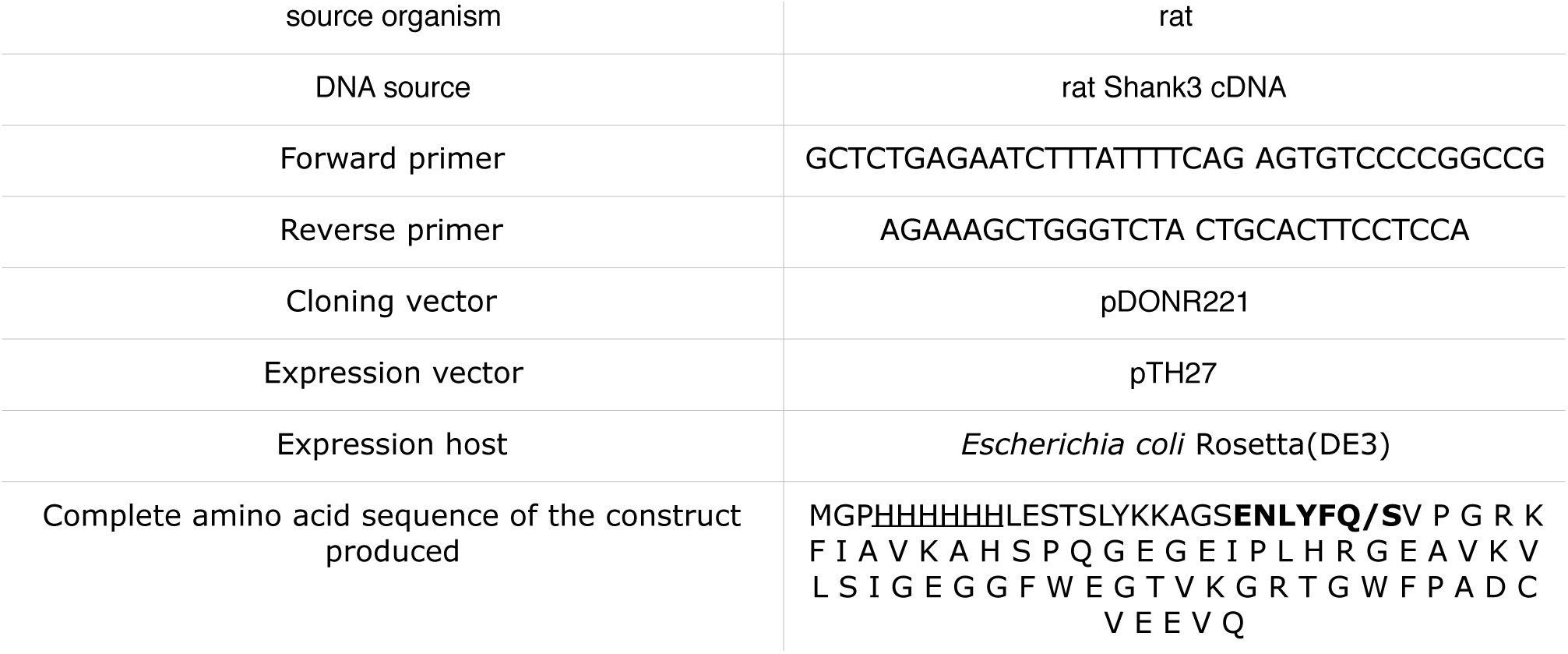
Macromolecule production information for Shank3-SH3. In the protein sequence, the His-tag is underlined, and the TEV cleavage site is in bold. The Shank3-SH3 domain has one extra Ser residue at its N terminus after cleavage.

Recombinant Shank3-SH3 was produced as a His-tagged version in *Escherichia coli* Rosetta(DE3) cells. The protein was expressed in 1 liter of autoinduction medium (Studier, 2005) at 310 K overnight. Cells were harvested and resuspended in 50 ml of lysis buffer (50 mM phosphate buffer, 300 mM NaCl, 10 mM imidazole, pH 7.0) and lysed with 10-15 cycles of sonication for 10 s each using a Branson sonicator and a 1/2" tapped horn. Cell debris was removed by centrifugation at 15000 rpm (27000 g) at 277 K for 40 min. The supernatant, containing the soluble Shank3-SH3 protein, was loaded onto a gravity flow Ni-NTA column. Then, the matrix was washed 3 times with 10 ml of lysis buffer, and the bound protein was eluted with elution buffer (50 mM phosphate buffer, 300 mM NaCl, 500 mM imidazole, pH 7.0).

The eluted protein was dialysed overnight against dialysis buffer (50 mM HEPES, 150mM NaCl, 1mM DTT, pH 7.0), and simultaneously, the affinity tag was cleaved using TEV protease (van den Berg *et al.*, 2006). The cleaved protein was then passed again through a Ni-NTA column to remove TEV protease, the His tag, and uncut protein. The protein was concentrated using 3K Amicon ultra centrifugal filters, and gel filtration was used as the final purification step prior to crystallisation screening. For gel filtration the same buffer as the dialysis buffer, but without DTT, was used. Shank-SH3 was run through a Superdex S75 16/60 column (GE Healthcare), and pure protein fractions were collected and concentrated to 20 mg/ml and used for crystallisation screening. Total yield of the protein was 6 mg (20 mg/ml in 300 u.1) from 1 liter of bacterial culture.

For crystallisation, Shank3-SH3 was concentrated to 20 mg/ml, and crystal screening was carried out in 96-well format using vapour diffusion in sitting drops. The optimised crystallisation conditions are given in Table 2.

**Table 2.** ***Crystallisation of Shank3-SH3***.

| Plate type | TTP 3-drop 96-well plate |
| --- | --- |
| Temperature (K) | 293 |
| Protein concentration | 20 mg/ml |
| Buffer composition of protein solution | 50 mM HEPES (pH 7.0), 150 mM NaCl |
| Composition of reservoir solution | 0.1 M bis-tris (pH 6.5), 32.5% PEG3350 |
| Volume and ratio of drop | 450 nl, drop ratio 2:1 (protein:buffer) |
| Volume of reservoir | 50 μl |

Prior to X-ray diffraction data collection, the crystals were cryo-protected (0.1 M bis-tris pH 6.5, 32.5% PEG3350, 20% PEG400) and rapidly cooled in liquid nitrogen. Data from several crystals of Shank3-SH3 were collected on beamline ID23-1 at ESRF, Grenoble, France. The data were processed with XDS (Kabsch, 1988, 2010). Data intensity distributions, as seen for example using phenix.xtriage (Zwart *et al.*, 2005), indicated all the crystals were nearly perfectly twinned, with the likely true space group being P2_1_ - with a fortuitous β angle of 90.05°. The estimated twin fractions for the operator h,-k,-l ranged between 36-46% for the different crystals. On the beamline setting, not all high-resolution data available for the best crystals could be collected due to geometric restraints. Data processing statistics for the best crystal that was tested, diffracting to the highest resolution and having the lowest estimated twin fraction, are shown in Table 3.

**Table 3.** Data collection and processing. Values for the outer shell are given in parentheses.

|  |  |
| --- | --- |
| Diffraction source | ESRF ID23-1 |
| Wavelength (Å) | 0.977 |
| Temperature (K) | 100 |
| Detector | Pilatus 6M |
| Crystal-detector distance (mm) | 121.56 |
| Rotation range per image (°) | 0.15 |
| Total rotation range (°) | 210 |
| Exposure time per image (s) | 0.037 |
| Space group | P2<sub>1</sub> |
| *a*, *b*, *c* (Å) | 29.74, 52.91, 31.66 |
| $\alpha$, $\beta$, $\gamma$ (°) | 90.00, 90.05, 90.00 |
| twinning operator (estimated twin fraction) | h,-k,-l (36%) |
| Resolution range (Å) | 50-0.87 (0.89-0.87) |
| Total No. of reflections | 229188 (1375) |
| No. of unique reflections | 64681 (698) |
| Completeness (%) | 81.0 (11.8) <sup>#</sup> |
| Redundancy | 3.5 (2.0) |
| $\langle I/\sigma(I) \rangle$ | 20.9 (2.1) |
| R<sub>meas</sub> | 0.043 (0.524) |
| Overall *B* factor from Wilson plot (Å<sup>2</sup>) | 9.2 |
| CC<sub>1/2</sub> (%) | 99.8 (75.7) |

<sup>#</sup> The low completeness at high resolution results from geometrical issues during data collection. For example, between 1.01-0.97 Å, the data are 90.0% complete, with $\langle I/\sigma(I) \rangle = 10.1$ and R<sub>meas</sub> = 0.122.

^#^ The low completeness at high resolution results from geometrical issues during data collection. For example, between 1.01-0.97 Å, the data are 90.0% complete, with 〈I/σ(I)〉 = 10.1 and R_meas_ = 0.122.

## Results

In order to obtain a better understanding at the structural level on the Shank family SH3 domains, we produced and crystallised the SH3 domain of rat Shank3. No specific binding partners have yet been confirmed at the molecular level for any of the Shank SH3 domains, despite extensive literature on the Shank proteins and their interactions. The amino acid sequence of the Shank3 SH3 domain is identical between mouse, rat, and humans, and SH3 domains from different Shank proteins are highly similar. Hence, it is likely that the SH3 domains of different Shank family members share similar properties.

The Shank3 SH3 domain could be produced recombinantly at high yields and purity (Fig. 1), and crystals of Shank3-SH3 were readily obtained using standard vapour diffusion methods (Fig. 2).

**Figure 1.**
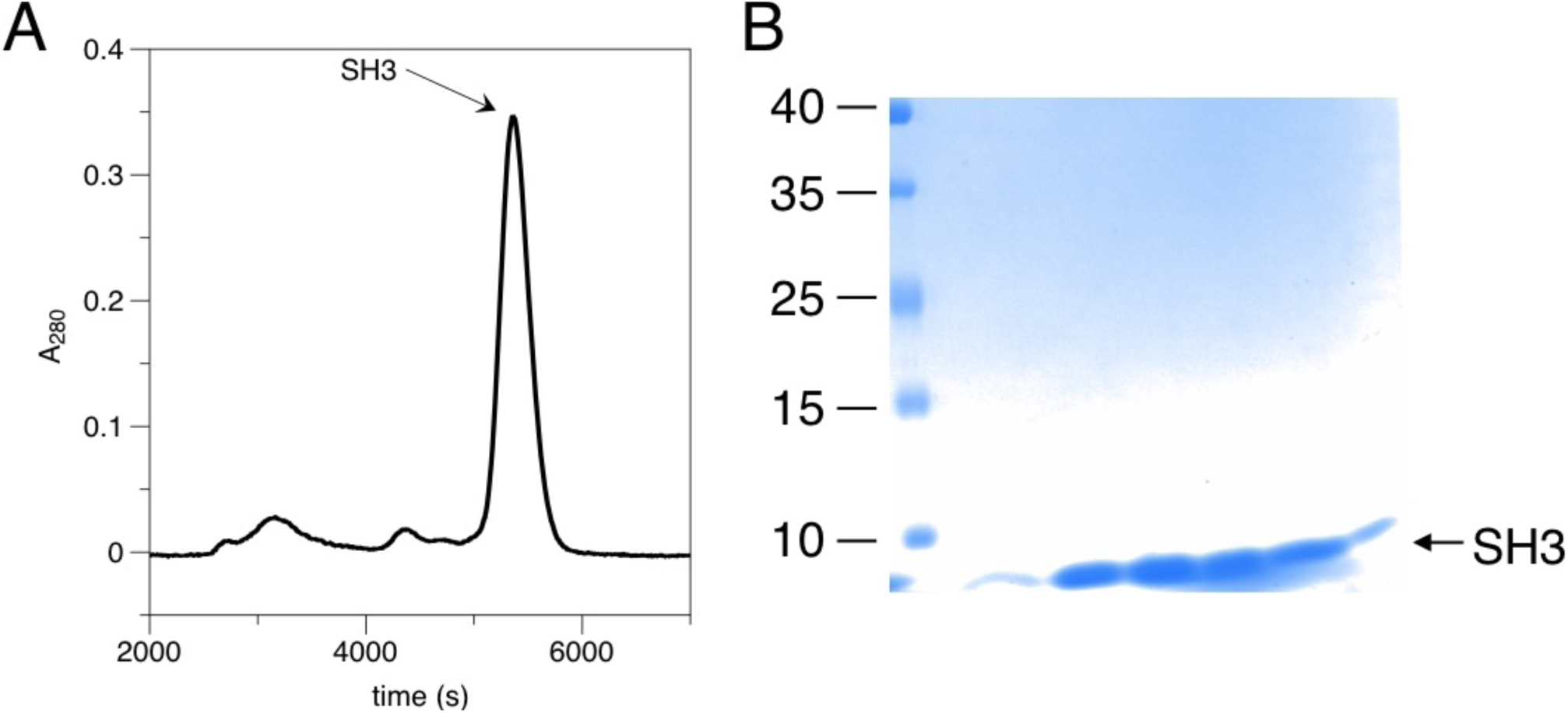
Purification of Shank3-SH3. A. Gel filtration of TEV-cleaved Shank3-SH3. The main peak corresponding to monomeric cleaved protein was collected. B. SDS-PAGE analysis of the gel filtration peak fractions. The position of the SH3 domain is indicated in both panels.

**Figure 2.**
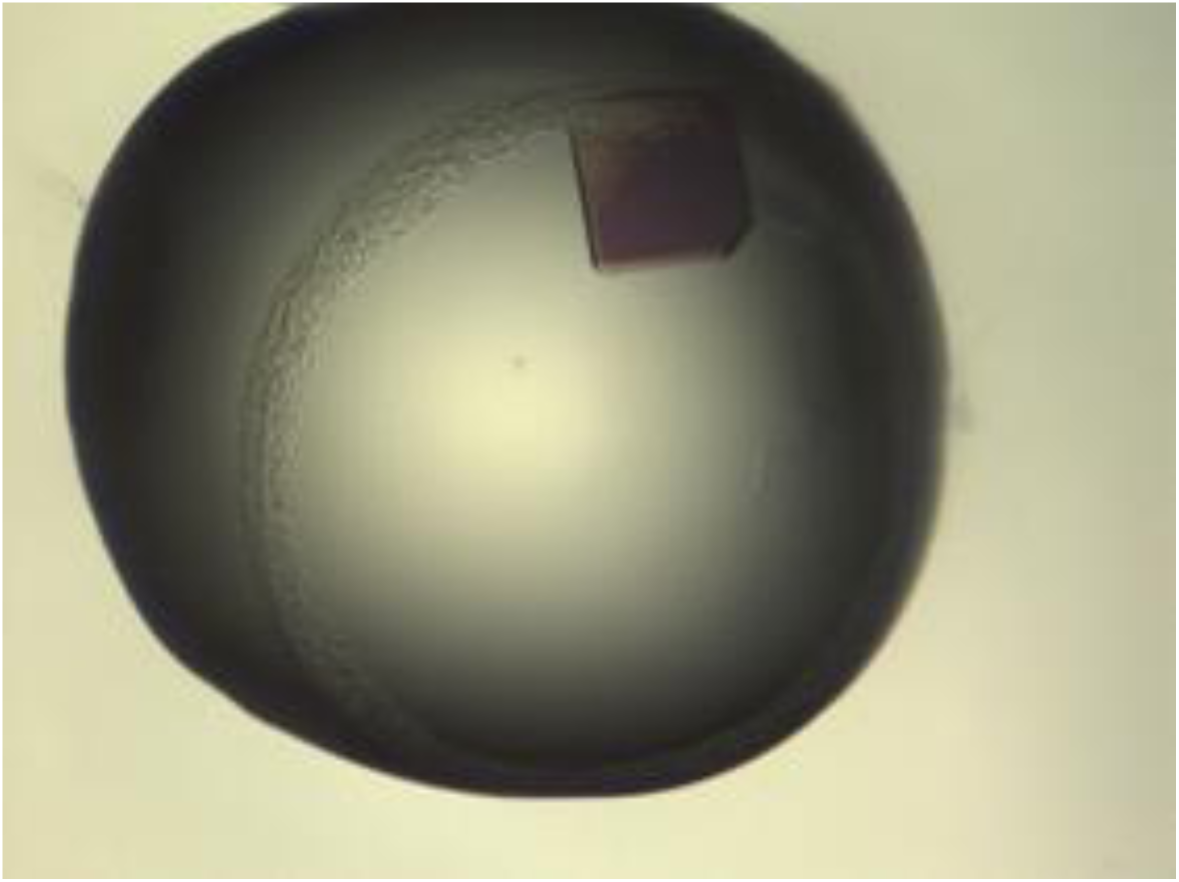
A typical single crystal of Shank3-SH3. The dimensions of the crystal were approx. 450 × 430 × 50 μm^3^.

The crystal used for the dataset described here was grown in 0.1 M bis-tris, 32.5% PEG3350. Diffraction could be observed to a resolution of at least 0.87 Å (Fig. 3).

**Figure 3.**
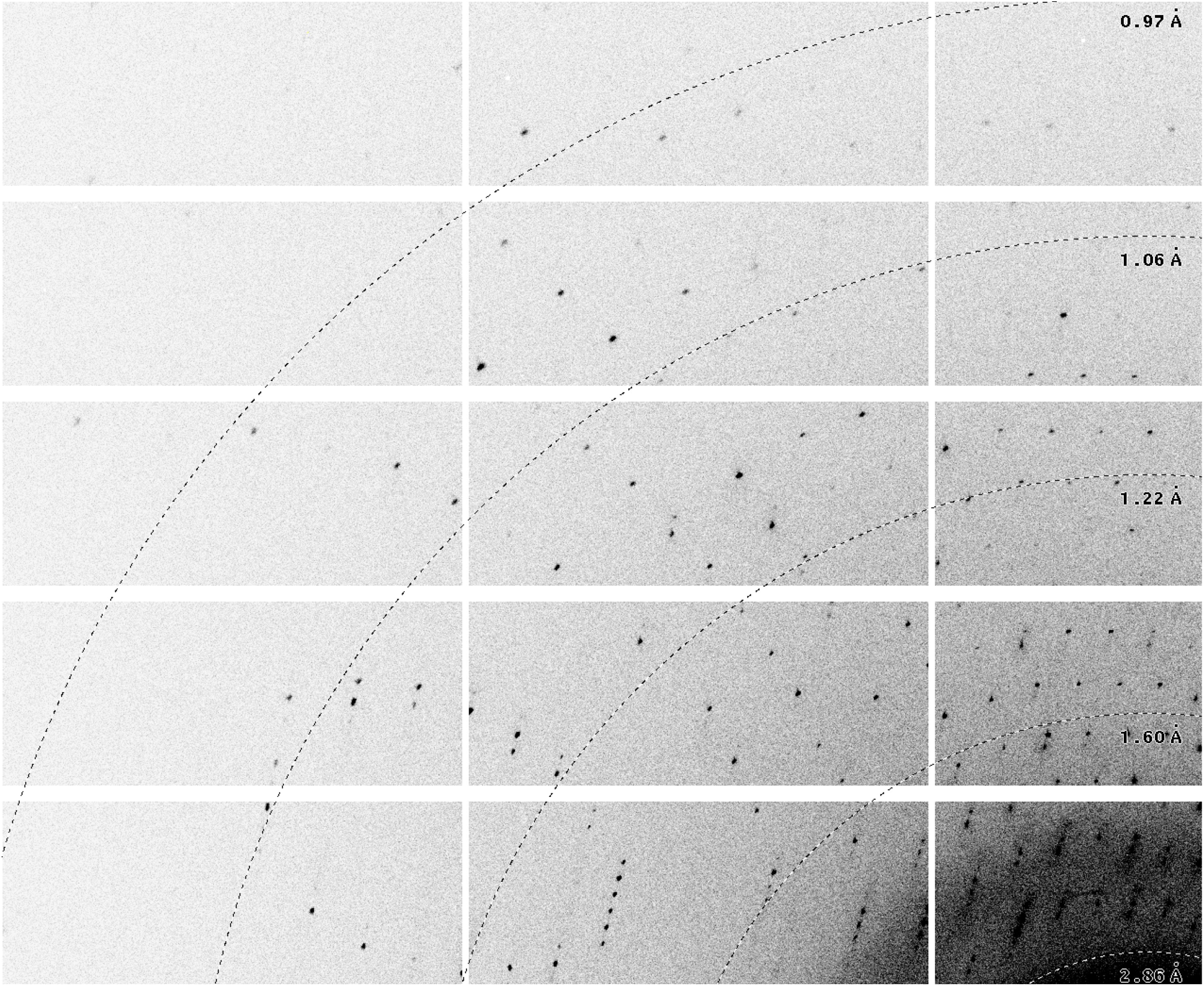
One corner of a diffraction image from the Shank3-SH3 domain crystal, collected on the ESRF beamline ID23-1. The resolution at the detector edge is 0.98 Å and in the corner (top left) 0.87 Å.

Initially, an orthorhombic lattice was selected by automatic processing routines at the beamline, but careful investigation of the intensity distributions indicated a high level of twinning with the operator h,-k,-l, and hence, all data were reprocessed in a monoclinic setting; systematic absences suggested space group P2_1_. The lowest estimated twinning fraction observed between different crystals was 36%. Thus, the crystals are not necessarily perfectly twinned, which will increase the chances of successful structure solution and refinement in the future.

The processing statistics (Table 3) indicate that the full diffraction capacity of the crystal could not be even nearly exploited during the data collection. It is expected that, using an optimised synchrotron beamline setting, we can collect even higher-resolution, complete data for Shank3-SH3 in the near future. This will eventually allow to pinpoint fine details of protein structure using the Shank3-SH3 domain as a tool. The more crucial step before this, however, will be the actual structure solution.

SH3 domains are small proteins containing β strands and flexible loops, and their structures are often difficult to solve by molecular replacement. The closest homologues to Shank3-SH3 in the PDB present sequence identities of ≈25% to Shank3, and presumably, experimental phasing will have to be employed to solve the structure. Such low sequence homology for a small domain of <60 residues dictates that molecular replacement is unlikely to work. Initial molecular replacement trials, using various SH3 domains from the PDB, have all failed (data not shown). On the other hand, the strong twinning component in the crystals is further likely to make structure solution and refinement complicated, even at sub-atomic resolution. Given all the caveats above, once the structure can be solved, it will provide a major increase in the understanding of Shank structure and interactions.

## Acknowledgements

This work has been supported by the Sigrid Jusélius Foundation (Finland), the Emil Aaltonen Foundation (Finland), and the Academy of Finland. The use of computing facilities at the Hamburg Centre for Bioinformatics, as well as beamtime and support at the ESRF, are gratefully acknowledged. We specifically wish to thank Juha Kallio for data collection at ESRF.

## References

Boeckers, T. M., Bockmann, J., Kreutz, M. R. & Gundelfinger, E. D. (2002). J Neurochem 81, 903–910.

Boeckers, T. M., Kreutz, M. R., Winter, C, Zuschratter, W., Smalla, K. H., Sanmarti-Vila, L, Wex, H., Langnaese, K., Bockmann, J., Garner, C. C. & Gundelfinger, E. D. (1999). J Neurosci 19, 6506–6518.

Durand, C. M., Betancur, C., Boeckers, T. M., Bockmann, J., Chaste, P., Fauchereau, F., Nygren, G., Rastam, M., Gillberg, I. C., Anckarsater, H., Sponheim, E., Goubran-Botros, H., Delorme, R., Chabane, N., Mouren-Simeoni, M. C., de Mas, P., Bieth, E., Roge, B., Heron, D., Burglen, L, Gillberg, C., Leboyer, M. & Bourgeron, T. (2007). Nat Genet 39, 25–27.

Grabrucker, S., Proepper, C., Mangus, K., Eckert, M., Chhabra, R., Schmeisser, M. J., Boeckers, T. M. & Grabrucker, A. M. (2014). Exp Neurol 253, 126–137.

Grubb, D. R., Iliades, P., Cooley, N., Yu, Y. L., Luo, J., Filtz, T. M. & Woodcock, E. A. (2011). FASEB J 25, 1040–1047.

Hammarström, M., Woestenenk, E. A., Hellgren, N., Hard, T. & Berglund, H. (2006). J Struct Funct Genomics 7, 1–14.

Han, K., Holder, J. L. J., Schaaf, C. P., Lu, H., Chen, H., Kang, H., Tang, J., Wu, Z., Hao, S., Cheung, S. W., Yu, P., Sun, H., Breman, A. M., Patel, A., Lu, H. C. & Zoghbi, H. Y. (2013). Nature 503, 72–77.

Kabsch, W. (2010). Acta Cryst D 66, 125–132.

Kabsch, W. (1988). J. Appl. Cryst. 21, 67–72.

Naisbitt, S., Kim, E., Tu, J. C., Xiao, B., Sala, C., Valtschanoff, J., Weinberg, R. J., Worley, P. F. & Sheng, M. (1999). Neuron 23, 569–582.

Sheng, M. & Kim, E. (2000). J Cell Sci 113, 1851–1856.

Studier, F.W. (2005). Protein Expr Purif 41, 207–234.

Tu, J. C., Xiao, B., Naisbitt, S., Yuan, J. P., Petralia, R. S., Brakeman, P., Doan, A., Aakalu, V. K., Lanahan, A. A., Sheng, M. & Worley, P. F. (1999). Neuron 23, 583–592.

van den Berg, S., Lofdahl, P. A., Hard, T. & Berglund, H. (2006). J Biotechnol 121, 291–298.

Wilson, H. L., Crolla, J. A., Walker, D., Artifoni, L., Dallapiccola, B., Takano, T., Vasudevan, P., Huang, S., Maloney, V., Yobb, T., Quarrell, O. & McDermid, H. E. (2008). Eur J Hum Genet 16, 1301–1310.

Zitzer, H., Honck, H. H., Bachner, D., Richter, D. & Kreienkamp, H. J. (1999). J Biol Chem 274, 32997–33001.

Zwart, P. H., Grosse-Kunstleve, R. W. & Adams, P. D. (2005). CCP4 Newsletter 42, 10.

